# Strong phylogenetic and ecological effects on host competency for avian influenza in Australian wild birds

**DOI:** 10.1101/2022.02.14.480463

**Authors:** Michelle Wille, Simeon Lisovski, David Roshier, Marta Ferenczi, Bethany J. Hoye, Trent Leen, Simone Warner, Ron A. M. Fouchier, Aeron C. Hurt, Edward C. Holmes, Marcel Klaassen

## Abstract

Host susceptibility to parasites is mediated by intrinsic and external factors such as genetics, age or season. While key features have been revealed for avian influenza A virus (AIV) in waterfowl of the Northern Hemisphere, the role of host phylogeny has received limited attention. Herein, we analysed 12339 oropharyngeal and cloacal swabs and 10826 serum samples collected over 11 years from wild birds in Australia. As well as describing species-level differences in prevalence and seroprevalence, we reveal that host phylogeny is a key driver in susceptibility. We confirm the role of age in AIV seroprevalence and viral prevalence. Seasonality effects appear less pronounced than in the Northern Hemisphere, while annual variations are potentially linked to El Niño– Southern Oscillation. Taken together, our study provides new insights into evolutionary ecology of AIV in its avian hosts, defining distinctive processes on the continent of Australia and expanding our understanding of AIV globally.

## Introduction

Wild birds are believed to be the reservoir for most influenza A viruses and have been detected across >100 avian species (Olsen *et al*. 2006). Avian influenza viruses (AIV) are predominately low pathogenic with limited signs of disease (Kuiken 2013). However, following spill-over into poultry, AIV may become highly pathogenic resulting in morbidity and mortality, thus causing substantial economic losses (Lycett *et al*. 2019), (Stamoulis 2017). There is also continued concern about zoonotic transmission of AIVs from poultry against the background of a continuously growing global poultry market (Naguib *et al*. 2019; Nunez & Ross 2019). A hallmark of this growing problem is spillback of highly pathogenic AIV into wild birds which results in mass mortality events in wild birds and the global spread of these viruses (Ramey *et al*. 2022).

Through intensive surveillance, members of the avian order *Anseriformes*, notably the family *Anatidae* (ducks, geese and swans), and to a lesser extent *Charadriiformes* (shorebirds and gulls) with emphasis on the family *Scolopacidae* (sandpipers), have been identified as key reservoirs of low pathogenic AIV (Olsen *et al*. 2006). Within these taxa, there appears to be significant heterogeneity in susceptibility, prevalence, viral diversity and host response to AIV across sampled host species (Olsen *et al*. 2006). Indeed, ducks of the genus *Anas* have generally been reported to have high prevalence and diversity of AIV subtypes (Olsen *et al*. 2006). This has led to an overrepresentation of key host taxa, including *Anas* ducks, in research systems.

In light of this bias, it is important to recognize that our current understanding of AIV ecology is described from a duck-centric, temperate and Northern Hemisphere perspective. Indeed, the ecology of AIV in *Anas* ducks, particularly Mallard (refer to Table S1 for scientific names) has been intensively interrogated at key sampling sites in Europe (Latorre-Margalef *et al*. 2014; van Dijk *et al*. 2014) and North America (Papp *et al*. 2017; Ramey & Reeves 2020). In these locations, AIV prevalence is highly seasonal, with a peak of 20-30% in the autumn as a result of recruitment of immunologically naïve individuals in the population following breeding, and migration-related aggregation of birds (Latorre-Margalef *et al*. 2014; van Dijk *et al*. 2014). However, a continental-scale study of AIV dynamics across North America demonstrated that infection dynamics may vary geographically due to differences in climate, seasonality and host ecology (Lisovski *et al*. 2017), with low-latitude environments having lower AIV prevalence with limited seasonal variation (Lisovski *et al*. 2017; Diskin *et al*. 2020). Studies in Africa reinforce these findings, with a detectable prevalence peak associated with the arrival of waterfowl migrants (Gaidet *et al*. 2012a; Gaidet 2016), rather than an association with season. Data from Australia have shown low prevalence in general and no consistent seasonal pattern (Hansbro *et al*. 2010; Grillo *et al*. 2015). Rather, profound inter-annual variation in the timing and quantity of rainfall, which is strongly linked with El Niño–Southern Oscillation (ENSO) and Indian Ocean Dipole (IOD), drives duck population breeding ecology and therewith AIV dynamics on this continent (Halse & Jaensch 1989; Norman & Nicholls 1991; Briggs 1992; Ferenczi *et al*. 2016; Stuecker *et al*. 2017). Beyond *Anseriformes*, AIV dynamics and ecology in *Scolopacidae* is unclear, with the exception of shorebirds sampled in Delaware Bay, USA (Maxted *et al*. 2016). Globally, the reported prevalence of AIV in shorebirds is low, and beyond Delaware Bay sampling is generally haphazard (Hanson *et al*. 2008; Winker *et al*. 2008; Gaidet *et al*. 2012b).

Taken together, we have a biased understanding of AIV ecology, with a strong focus on *Anas* ducks as reservoirs, and only a limited appreciation of geographic variations in these dynamics. Herein, we aim to address a number of key questions arising from this bias. First, to reveal the extent to which host species exhibit phylogenetically conserved patterns of susceptibility, which has recently been shown to be a critical aspect in patterns of host susceptibility (Longdon *et al*. 2011; Barrow *et al*. 2019). While species-level differences in prevalence are often reported in AIV studies, the role of phylogeny as a driver of these differences has never been incorporated, at either high (*i*.*e*. among avian orders) nor low (*e*.*g*. within families) levels of classification. Second, while controlling for these phylogenetic and species effects, we revisit the effects of age, season and eco-region as key ecological factors known to play a role in AIV prevalence, particularly in a geographic and climatic region that has seen limited research into AIV ecology. We address these questions based on the analysis of >10,000 samples collected over 11 years and across 76 species and seven avian orders, allowing for both a broad and an in-depth phylogenetic comparison across a wide host landscape for this virus. Critically, we leverage both virological and serological data into our framework. While virological data is central to understanding active infection, it may be deficient when sampling sporadically or without prior information on timing, age or species to target. As such, the addition of serological data allows us to garner a more complete picture of AIV dynamics on this unique continent.

## Methods

### Ethics Statement

All required ethics approvals and Australian state and territory permits were obtained prior to the catching and sampling of birds contained in this study (Detailed Ethics Statement in the Supplement).

### Sample collection

Samples were collected between November 2010 and March 2021. Three main catching techniques were employed. Baited funnel walk-in traps were deployed on land or in shallow water allowing for foraging by dabbling ducks. Traps baited with seeds were set at dawn and operated during the day and left open (birds could enter and leave the traps freely) during the night. Cannon nets to capture roosting ducks and shorebirds were operated during the day. To capture waterbirds at night, mist nets were erected on poles above the water surface. Small songbirds, doves and parrots were caught during the day using mist nets. All trapping techniques were used in areas of high bird activity (Whitworth 2007). Commencing in June 2016, hunted ducks were sampled within 12 hours of collection.

Both oropharyngeal and cloacal samples were collected from each individual bird using a sterile tipped applicator and placed into virus transport media (VTM, brain heart infusion [BHI] broth-based medium [Oxoid] with 0.3 mg/ml penicillin, 5 mg/ml streptomycin, 0.1 mg/ml gentamicin, and 2.5 g/ml, amphotericin B). Initially oropharyngeal and cloacal samples were placed in separate vials, while starting March 2014 these samples were pooled into a single vial containing VTM. Following collection, samples were kept cool (4-8ºC) for up to a week prior to being stored at -80ºC, or were stored in liquid nitrogen (−196 ºC) until they could be placed in a -º80C freezer.

Blood samples were collected from each bird, except for the hunted ducks. Up to 200µl was collected from the brachial vein using the Microvette capillary system for serum collection (Sarstedt). Occasionally blood samples were collected from the medial metatarsal vein of ducks. Following collection, samples were kept cool (4-8ºC) and 7-14 hours following collection were centrifuged and sera collected and stored at -20ºC.

### Sample screening for AIV infection

Samples collected in 2010 were assayed by the Australian Centre for Disease Preparedness as per (Curran *et al*. 2014). Between 2011-2015, samples were assayed at the Victorian Department of Economic Development, Biosciences Research Division. Briefly, RNA was extracted using the MagMax 96 Viral Isolation Kit (Ambio, Thermo Fisher Scientific) using the Kingfisher Flex platform (Thermo Fisher Scientific). RNA was assayed for a short fragment of the matrix gene (Fouchier *et al*. 2000). First, using the Superscript III Platinum ONE step qPCR Kit (Life Technologies, Thermo Fisher Scientific) with ROX, followed by a subsequent amplification and detection using the SYBR Green mastermix (Life Technologies, Thermo Fisher Scientific). Starting in 2015, all samples were assayed at the WHO Collaborating Centre for Reference and Research on Influenza (WHOCCRI). RNA was extracted using the NucleoMag Vet Kit (Scientifix) on the Kingfisher Flex System. Extracted RNA was subsequently assayed for a short fragment of the influenza A matrix gene (Spackman *et al*. 2002) using the SensiFAST Probe Lo-Rox qPCR Kit (Bioline). A cycle threshold (Ct) cut-off of 40 was used.

### Detection of anti-NP antibodies

Samples collected in 2010 were analysed as per (Curran *et al*. 2015). Starting in 2011 serum samples were assayed using the Multi Screen Avian Influenza Virus Antibody Test Kit (IDEXX, Hoppendorf, The Netherlands) following manufacturer’s instructions where an S/N value <0.5 was considered positive. Analyses prior to May 2015 were carried out in duplicate.

### Data analysis

For oropharyngeal and cloacal swab samples collected separately, we considered an individual bird positive if either the oral or cloacal sample were positive and merged into a single entry.

All analyses were conducted using R version 4.0.3 integrated into RStudio version 1.3.1093.

We used phylogenetic generalised linear mixed effect models to investigate the simultaneous effects of species as a random variable and fixed-effect, explanatory variables age, eco-region, season, and year on AIV prevalence and seroprevalence. For species we evaluated both the phylogenetic species effect, which evaluates the contribution of shared evolutionary history among species (*e*.*g*. genetic factors; termed “phylogenetic effect”) as well as the species effect independent of the phylogenetic relationship between species (*e*.*g*. ecological factors; termed “species effect”). Bird age was presented in two categories: juvenile (*i*.*e*. hatch-year) or adult. Three eco-regions, *i*.*e*. temperate, arid, and tropical, were used based on the 2012 Interim Biogeographic Regionalisation for Australia version 7 (https://www.environment.gov.au/land/nrs/science/ibra#ibra). Season was divided into summer (September-February) and winter (March-August). For migratory shorebirds, summer coincides with the arrival of birds from the breeding grounds followed by their primary moult. Winter includes the period of pre-migratory preparation prior to departure for northern hemisphere breeding grounds. This behaviour applies to birds in their 2nd year and older; birds in their first year remain in Australia for the southern hemisphere winter.

Species with fewer than 50 samples were excluded from the analyses. To evaluate phylogenetic and species effects across higher and lower levels of classification (*i*.*e*. comparing species across multiple orders versus a comparison of species within families), we ran analyses on three sets of taxa. First, a set containing all species sampled. Second, a subset of this first group with only species belonging to the family *Anatidae* and, third, only species belonging to the family *Scolopacidae*. For the latter two taxon sets, we removed year 2014, 2015 and 2016 and the tropical eco-region for Anatidae, and the arid eco-region for *Scolopacidae*, due to low sample sizes.

The analyses were conducted using Bayesian generalized (non-) linear multivariate multilevel models using the brm() function within R package brms (Bayesian Regression Models using ‘Stan’), using family “Bernoulli” and default priors (Bürkner 2017; Bürkner 2018). We ran a series of candidate models for each of the three taxon sets and for both AIV and seroprevalence, *i*.*e*. 6 model sets with 10 models each, following (Barrow *et al*. 2019): (1) a model containing only an intercept, (2) a model containing an intercept plus the phylogenetic and species effects (3) the full model containing all fixed-effect explanatory variables as well as the phylogenetic and the species random effects, (4) the full model minus the phylogenetic effect, (5) the full model minus the species effect, (6) the full model without phylogenetic and species effects, (7) the reduced model, (8) the reduced model without the phylogenetic effect, (9) the reduced model without the species effect, and (10) the reduced model without the species and phylogenetic effects. The reduced models included only the predictors found to be important, *i*.*e*. their 95% credible intervals (CI) were non-overlapping with zero in the full models. Each model was run using four chains for 2,000 generations, retaining the final 1,000 for which we used a thinning factor of 5 to determine the posterior distributions of the parameters. Convergence was assessed using trace plots and scale reduction factors 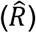, where we accepted convergence with 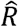 values less than 1.02, with a preference for values of less than 1.01. For each model the Widely Applicable Information Criterion (WAIC) was calculated using the *loo* package (Vehtari *et al*. 2017) and used to compare the 10 models within a set (Watanabe 2010). Models that had a ΔWAIC ≤ 2 (*i*.*e*. were within 2 WAIC units of the model within the set of 10 with the lowest WAIC) were considered to be models describing the data set best. Phylogenetic signal (λ) was computed following (Bürkner 2021), substituting σ^2^ with π^2^/3, since we are dealing with a Bernoulli distribution.

Host phylogenies used in the analyses were estimated using concatenations of RAG-1 (Recombination activating gene 1), CytB (Cytochrome B), COI (Cytochrome c oxidase I) and N2 (NADH dehydrogenase 2) genes available in GenBank. Concatenated sequences were aligned using the MAAFT algorithm (Katoh & Standley 2013) within Geneious R11, and a maximum likelihood tree was inferred incorporating the best fitting nucleotide substitution model in PhyML 3.0 (Guindon *et al*. 2010). Phylogenies were confirmed to conform to those previously determined, as described by (Baker *et al*. 2007; Gibson & Baker 2012; Sun *et al*. 2017).

## Results

### Sampling regime

Between 2010-2021, 10826 serum samples and 12339 swab samples (combined oropharangeal and cloacal) were collected in Australia. The data set comprises 11 orders, 25 families, and 75 species of Australian birds, although the majority of the samples were collected from members of the family *Anatidae* within the Anseriformes (3657 swabs and 2412 serum samples) and family *Scolopacidae* within the *Charadriiformes* (7622 swabs and 7520 serum samples) (Fig 1). Avian orders for which we had negligible sample numbers included the *Galliformes* (n=4), *Podicipediformes* (n=7) and *Suliformes* (n=3) (Table S1).

**Figure 1.**
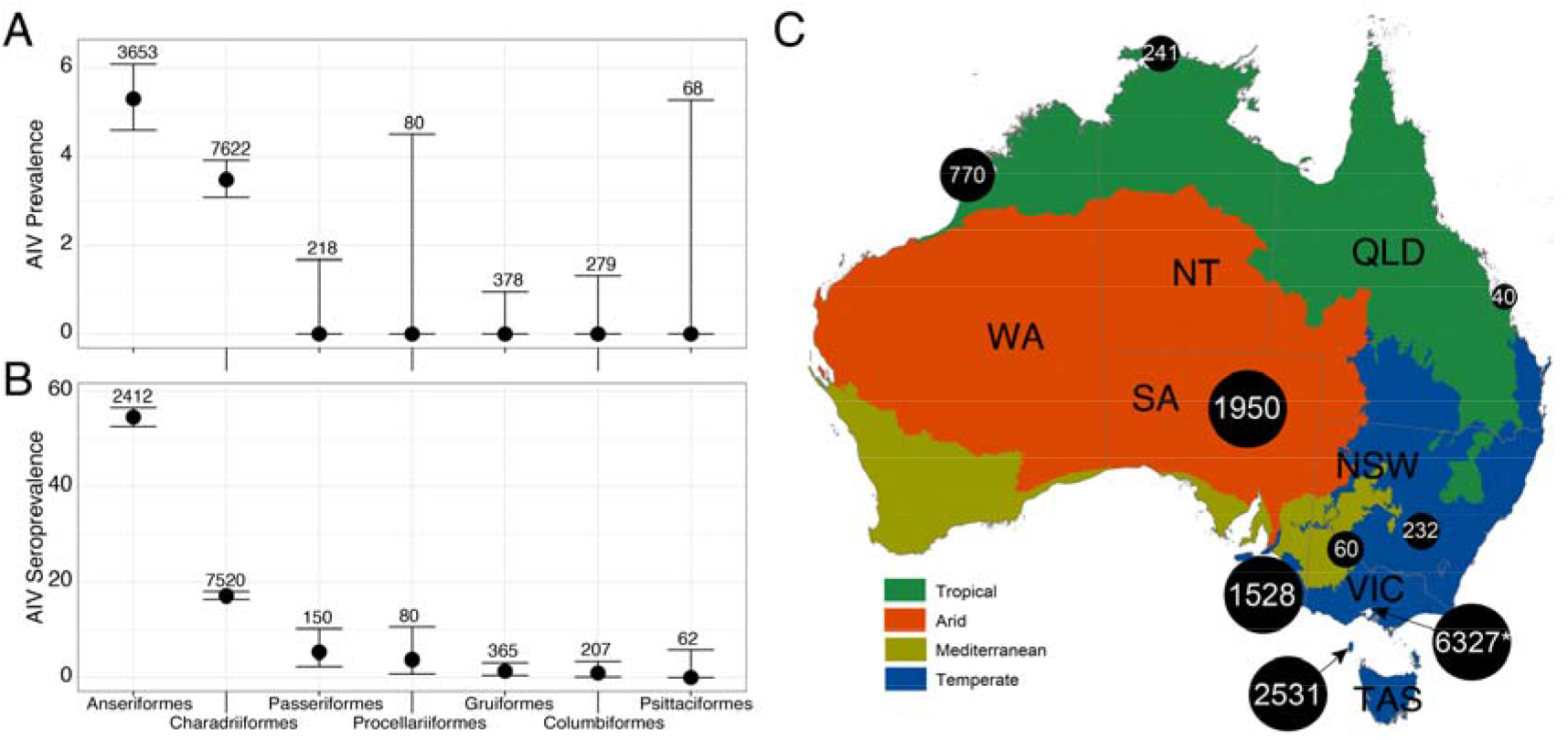
Sampling effort and virus prevalence across the study period. (A) Avian influenza viral prevalence based on qPCR of swab samples and (B) seroprevalence based upon a commercial anti-NP ELISA of serum samples. Points represent point estimates of percentage prevalence and bars are the 95% confidence interval. Numbers above each estimate represent the number of samples included. For both A and B, we excluded avian orders from which we had <10 samples collected throughout the entire study period (C) Map illustrating geographic sampling effort across Australia. Map colours refer to eco-regions and was generated from https://www.environment.gov.au/land/nrs/science/ibra/australias-ecoregions, and distributed under a Creative Commons Attribution 3.0 Australia License. Herein “Temperate” includes both temperate grasslands and forests, “Tropical” includes tropical and subtropical forests and grasslands, “Arid” includes deserts and xeric shrublands. Australian state names are TAS Tasmania, VIC Victoria, SA South Australia, NSW New South Wales, WA Western Australia, NT Northern Territory, QLD Queensland. Sampling location is indicated by a black circle, with size corresponding to the number of individuals sampled. Numbers within black circles refer to the number of individuals sampled; for some individuals we may have both swab and serum samples and for others only swab or serum samples. Those samples collected from Victoria, but not in study sites in and around Melbourne have been added to the Victorian count, and this is indicated by an asterisk. A detailed breakdown of species composition is presented in Table S1.

Overall, we found evidence of AIV infection and anti-AIV antibodies in *Anseriformes* (5.4% virus prevalence and 53% seroprevalence) and *Charadriiformes* (3.5% virus prevalence and 17% seroprevalence), with 4% virus prevalence and 17% seroprevalence in the *Scolopacidae*. This is in accord with our expectation that members of these two orders (and families) of birds comprise the main AIV reservoirs. While we failed to find active AIV infection, we did detect low level seropositivity in the *Passeriformes* (5.3%), *Procellariforms* 3.8%), *Gruiiformes* (1.4%) and *Columbiformes* (0.97%). We found no evidence of anti-AIV antibodies in any of the 62 *Psittaciformes* tested (Fig 1).

All orders were heavily targeted in temperate sites in the states of Victoria, Tasmania and South Australia, with the largest number of samples collected from the Western Treatment Plant, Victoria (38°00’S, 144°34’E) [2759 swabs, 2769 serum], Western Port Bay, Victoria (38°12’40”S, 145°22’48”E) [1365 swabs, 1092 serum], King Island, Tasmania (39°52’S, 143°59’E) [2522 swabs, 2313 serum] and Limestone Coast, South Australia (37°27’05”S 139°58’42”E) [746 swabs, 808 serum]. Between 2011-2014, samples were collected from the arid Innaminka Regional Reserve, South Australia (27°32’28”S, 140°35’47”E) [1936 swab and 1609 serum samples], and samples from three independent sampling expeditions were collected in tropical Northwestern Western Australia (17°58’10”S 122°19’23”E) [739 swab, 766 serum]. An additional 2272 swab and 1469 serum samples were collected from other sites in Australia (Fig 1).

### Phylogenetic and non-phylogenetic species effects are key determinants of host competence

Six different species within the *Anseriformes* were included in our analysis: Australian Shelduck, Australian Wood Duck, Pink-eared duck and three members of the genus *Anas*: Grey Teal, Chestnut Teal and Pacific Black Duck. While overall, seroprevalence was 53% and virus prevalence 5.4% for *Anseriformes*, Wood Duck had a significantly lower seroprevalence (2.8%) and viral prevalence (2.3%) compared to other species suggesting that it is a less competent AIV host (Fig 2).

**Figure 2.**
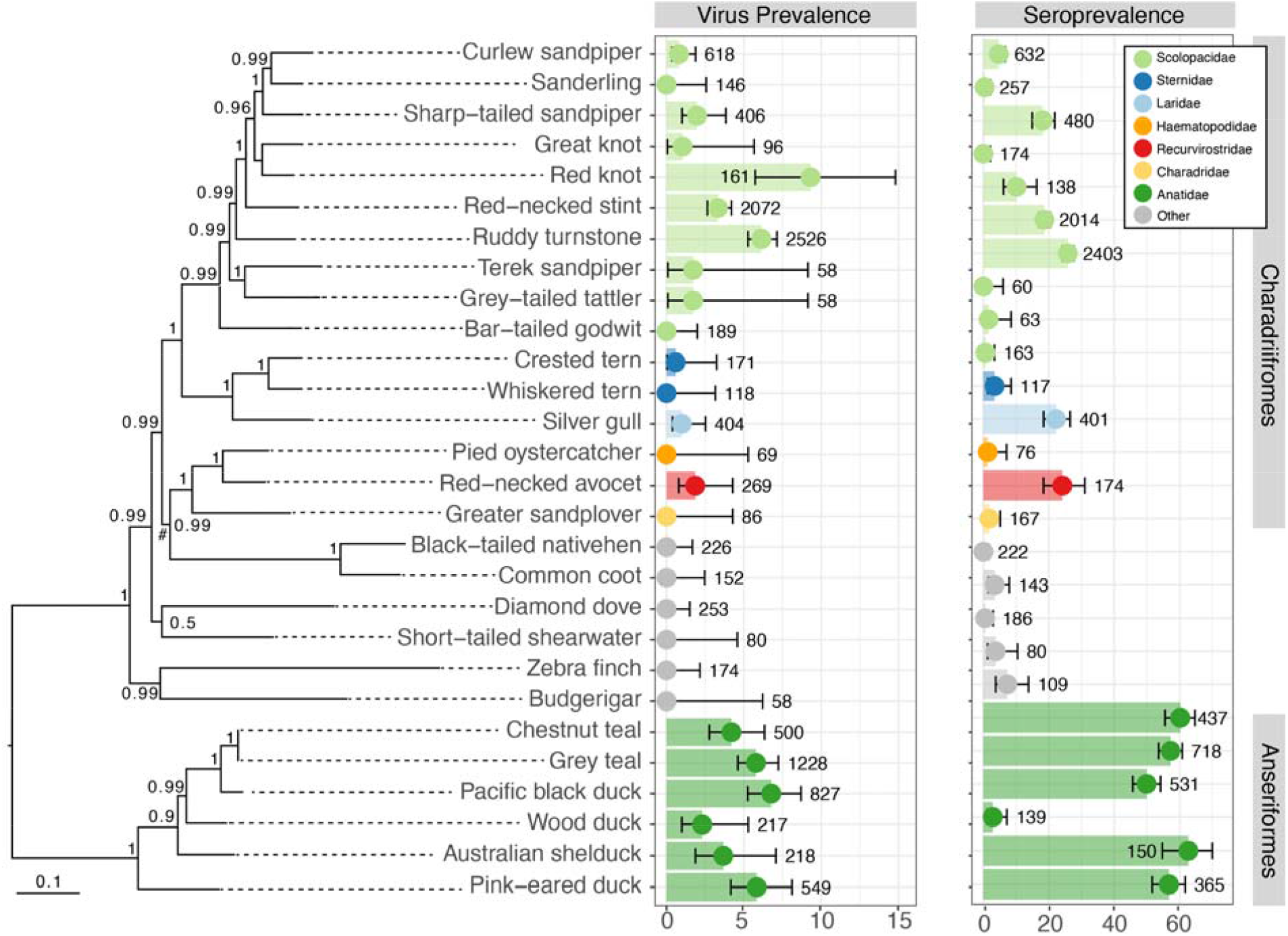
Prevalence and seroprevalence in Anseriformes and Charadriiformes. Host species are arranged taxonomically, according to maximum likelihood phylogenies based on concatenated mitochondrial and one nuclear marker. aBayes support values are presented at the node, and the scale bar indicates the number of substitutions per site. Hash/pound (#) refers to a node on the tree which does not conform to known phylogenetic relationships and we were unable to resolve this discrepancy based on available host genetic data in GenBank. Species from which >50 samples collected are included, and percentage prevalence and 95% confidence intervals are plotted. Colours refer to host families in the order Anseriformes and Charadriifromes, host families from other orders are in grey. Sample size is plotted adjacent to the point estimate. Seroprevalence refers to the percentage prevalence of anti-NP antibodies in collected serum samples. Virus prevalence refers to the percentage prevalence of AIV by use of qPCR.

For the second most important host order for AIV, the *Charadriiformes*, shorebird species where >50 samples were collected included Red-necked Avocet, Pied Oystercatcher, Greater Sand Plover and from the *Scolopacidae*: Bar-tailed Godwit, Terek Sandpiper, Grey-tailed Tattler, Ruddy Turnstone, Great Knot, Red Knot, Curlew Sandpiper, Sanderling, Sharp-tailed Sandpiper, and Red-necked Stint. There was marked heterogeneity in both seroprevalence and viral prevalence across shorebird species. For example, in the *Scolopacidae*, we found higher viral prevalence and seroprevalence (>10%) in Ruddy Turnstone, Red Knot, Sharp-tailed Sandpiper and Red-necked Stint, with very low or no evidence of antibodies in Bar-tailed Godwit, Great Knot, Curlew Sandpiper and Sanderling. The only other shorebird from which we detected AIV was Red-necked Avocet (Fig 2). In addition to shorebirds, the *Charadriiformes* also include gulls and terns, and herein we included Silver Gull, Whiskered Tern, and Crested Tern. Viral prevalence was low (< 1%) in all three species, although seroprevalence in Silver Gulls was 22.2% (Fig 2), suggesting our sampling regime to detect AIV infection was possibly inadequate or NP-antibodies in this species are particularly long-lived.

For other species for which we collected more than 50 samples, viral prevalence was 0% (*i*.*e*. no AIV positive samples), and AIV seroprevalence was negligible (∼3%, except for Budgerigar wherein none of the serum samples were positive for anti-AIV antibodies).

Across all 10 candidate models tested for each of the three avian taxon sets, the models that considered phylogeny and species were the best fit for both virus prevalence and seroprevalence (*i*.*e*. had a ΔWAIC ≤ 2; Table 1). Further, models that included all or a reduced set of explanatory variables, as compared to neither, greatly improved the performance of the candidate models in describing the three taxon sets across both virus prevalence and seroprevalence. As such, models including phylogenetic effects, species effects and other variables (such as age, eco-region, year and season) are required to adequately explain AIV variation.

**Table 1.** ΔWAIC values for all 10 candidate models, for both virus prevalence and seroprevalence in 3 different host taxon sets. Models that fit the data most satisfactorily (with a ΔWAIC ≤ 2) are emboldened. Models are ranked based on overall performance, starting with models that tended to perform best in describing virus prevalence and seroprevalences across all three taxon sets. Generally, models that included both random effects (phylogeny and species), as well as a full or reduced set of fixed-effect, explanatory variables (age, eco-region, season, year) performed best in explaining the variation across all taxon sets, for both virus prevalence and seroprevalence.

| Model Description | | | | | Response ΔWAIC for prevalence in | | | | | |
| --- | --- | --- | --- | --- | --- | --- | --- | --- | --- | --- |
| Predictors |  |  | Random Effects |  | All species |  | Anatidae |  | Scolopacidae |  |
| All | Reduced | None | Species | Phylogeny | Virus | Serology | Virus<sup>3</sup> | Serology<sup>3</sup> | Virus<sup>1,2</sup> | Serology<sup>2</sup> |
| X |  |  | X | X | 3.7 | **0.6** | **1.3** | **0.0** | **0.2** | **1.7** |
| X |  |  |  | X | 2.5 | **0.7** | **0.4** | **0.3** | **1.2** | 2.7 |
|  | X |  | X | X | **1.5** | **0.0** | 3.3 | 6.9 | **0.2** | **0.0** |
|  | X |  |  | X | **0.0** | **1.1** | 2.3 | 6.3 | **1.1** | **1.4** |
| X |  |  | X |  | 9.6 | **0.9** | **0.7** | **0.2** | **0.1** | **1.4** |
|  | X |  | X |  | 7.2 | **0.7** | 3.4 | 6.8 | **0.0** | **0.5** |
|  |  | X | X | X | 141.6 | 459.9 | 40.8 | 68.9 | 90.7 | 405.2 |
| X |  |  |  |  | 111.4 | 1529.3 | **0.0** | 135.2 | 32.0 | 396.3 |
|  | X |  |  |  | 140.0 | 1529.1 | 2.1 | 149.6 | 30.1 | 445.3 |
|  |  | X |  |  | 335.7 | 2367.3 | 44.0 | 242.4 | 169.8 | 830.2 |

<sup>1</sup> The models did not contain Year as an explanatory variable, given poor model convergence when included.
<sup>2</sup> The models did not contain the arid eco-region due to low sample size.
<sup>3</sup> The models did not contain years 2014, 2015 and 2019, or the tropical eco-region due to low sample size.

Considering all species, the phylogenetic signal, λ, which can vary between 0 (non-existent) to 1 (very strong) was generally strong in both viral prevalence (0.76) and seroprevalence (0.71; Table 2; Fig 3). In addition to all species, we analysed the phylogenetic effect at two lower taxonomic levels (within the *Anatidae* and *Scolopacidae*). Within these key host families, the phylogenetic signals remained significant, varying between 0.27 and 0.60 (Table 2; Fig 3). It is noteworthy that the phylogenetic effect at these lower (family) level comparisons was lower than when comparing species across the seven orders (Table 2). However, it still showed that within these two major AIV host groups, considerable variation in competence levels exist between species. These species differences with a phylogenetic origin are further augmented by non-phylogenetic species differences, potentially related to differences in ecology, as evidenced by significant species signals among the top-ranking models in all taxon sets for both virus prevalence and seroprevalence (Fig 3, Table 1).

**Table 2.**
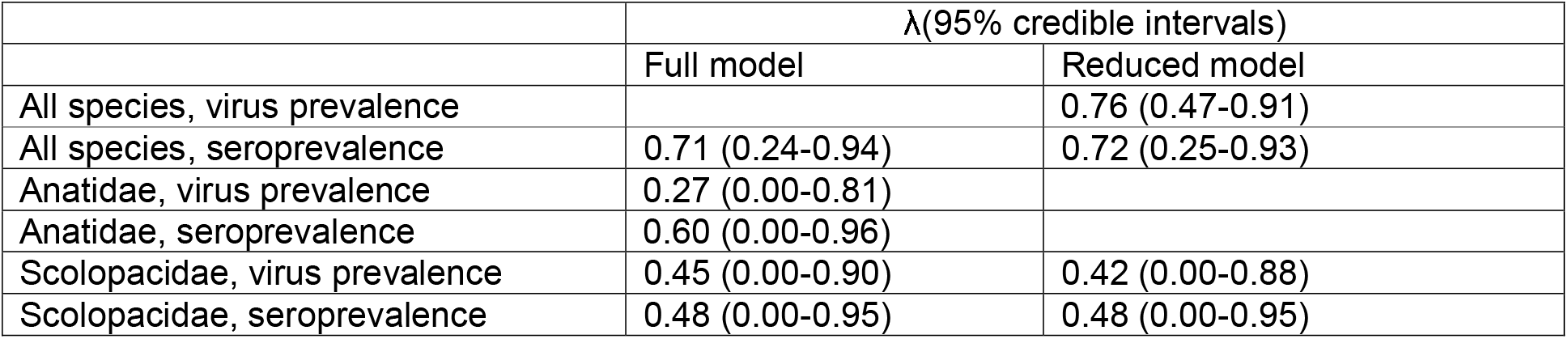
Phylogenetic signal (λ) estimates with 95% credible intervals for full and reduced models with both phylogeny and species as random effects. Phylogenetic signals are only indicated if the model had a ΔWAIC ≤ 2.

**Figure 3.**
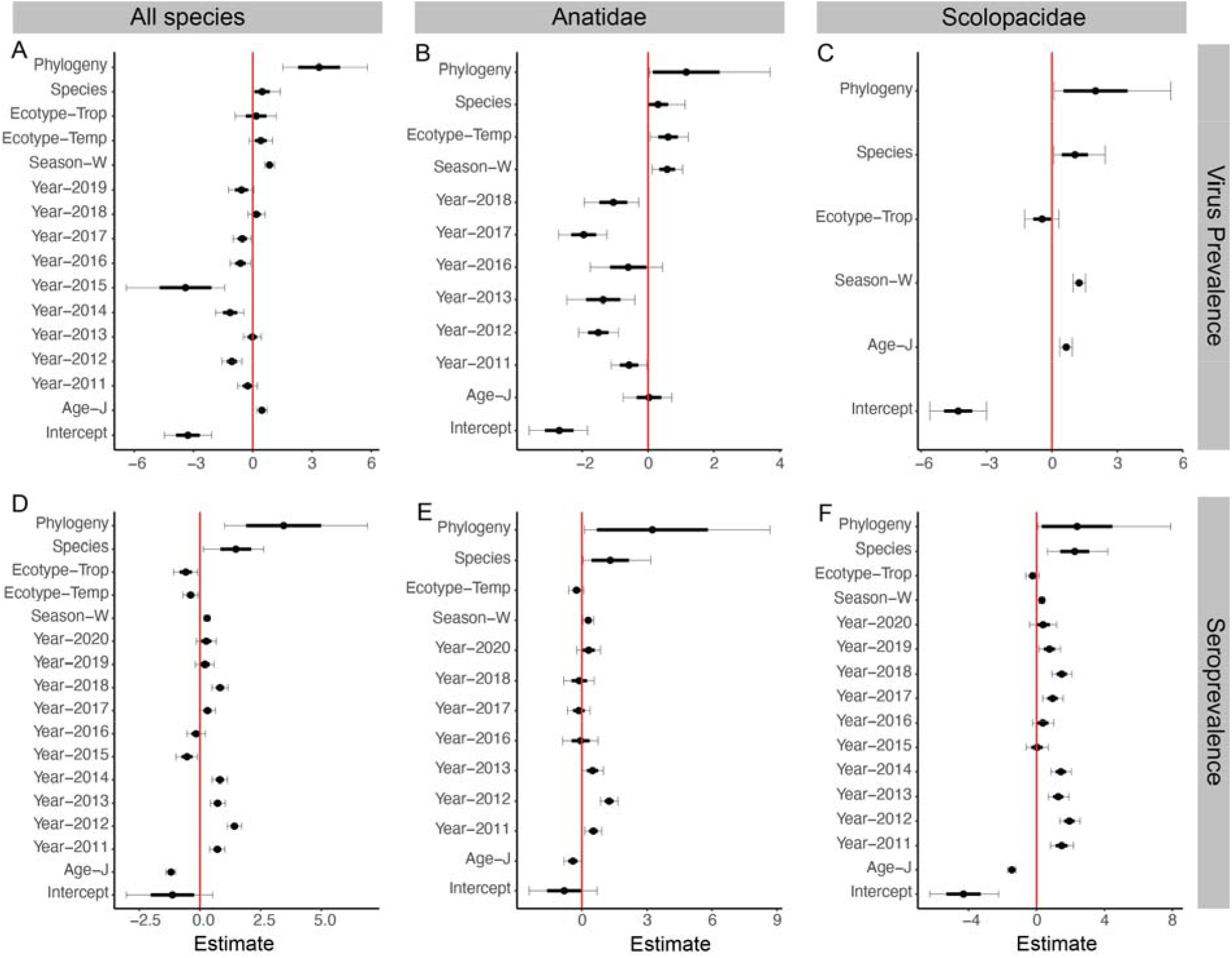
Posterior mean estimates with standard error (thick bars) and 95% credible intervals (capped thin bars) of predictors and random effects on (A, B, C) AIV prevalence and (D, E, F) seroprevalence for (A, D) all species with >50 samples, (B, E) Anatidae and (C, F) Scolopacidae for the full brms models. Parameters with intervals that do not overlap zero (indicated by a red line) are considered to have a significant influence on the response. Note that estimates are logits.

### Seroprevalence and viral prevalence have inverse relationships with bird age

Across the four explanatory variables investigated, age was an important predictor of virus prevalence and seroprevalence in the models covering all species with the *Scolopacidae*, and the *Anatidae* showing a similar tendency (Fig 3, Fig S1A). Across all species combined, juveniles had a 2.0% higher viral prevalence (95% CI 0.7─3.5%) and a 15.5% (−17.0─-13.7%) lower seroprevalence as compared to adults (where percentages are calculated from the logit estimates depicted in Fig 3). For the *Scolopacidae* only, these differences were 1.2% (0.5-2.0%) and -1.0% (−1.1 - -0.9%), for virus prevalence and seroprevalence, respectively. At a species level, significant differences in prevalence and seroprevalence between adults and juveniles were limited to species in which prevalence levels were also high for *Scolopacidae* (*i*.*e*. those species with a seroprevalence > 18%): Red-necked Stint, Ruddy Turnstone, and Sharp-tailed Sandpiper (Fig S1B). Trending in a similar direction, in *Anatidae* there was no significant age effect in virus prevalence (0.2%, 95% CI -3.2 – 5.8%) but there was in seroprevalence (−8.0%, -14.5 - 0.0%). At the species level within the *Anatidae*, there were no species where virus prevalence for juveniles were different from adults (Fig S1C). However, for both Pacific Black Duck and Pink-eared Ducks the seroprevalence estimates for juveniles were lower as compared to adults (Fig S1C). Unfortunately, sample size for juvenile *Anatidae* was generally low (<50, with the exception of Grey Teal for which we had 66 samples), which may have played a role in a limited age-dependant effect on prevalence and seroprevalence for this avian family. As prevalence for avian species not in the *Anseriformes* or *Charadriiformes* was negligible (0% virus prevalence, and 11% seroprevalence) no age-dependant patterns can be inferred for these orders.

### Season and year modulate AIV prevalence and seroprevalence

Although less pronounced than northern hemisphere studies, season significantly affected prevalence levels in our data. Across all three species groups, winter viral prevalence was significantly higher compared to summer viral prevalence estimates (where again percentages are calculated from the logit estimates depicted in Fig 3; all-species: 4.4%, 2.8 – 6.5; *Anatidae*: 4.4%, 0.8 – 9.8; *Scolopacidae* 3.2%, 2.1 – 4.6). Similarly, the same pattern was found in seroprevalence across all three species groups (all-species: 6.1%, 3.3 – 8.7; *Anatidae*: 6.5%, 1.1 – 12.7; *Scolopacidae* 0.5%, 0.2 – 0.8). In the case of *Scolopacidae* summer includes the arrival of birds from their Arctic breeding grounds while winter includes birds sampled during the pre-migration phase..

In all three taxon groups, sampling year drove significant variation in virus prevalence, except for *Scolopacidae*, wherein the model including year did not converge. Given strong year effects for both the *Anatidae* (virus prevalence and seroprevalence) and *Scolopacide* (seroprevalence only), it is unsurprising that there was also a strong year effect in the all-species models. Based on the findings of (Ferenczi *et al*. 2016), we compared annual rainfall across the Murray-Darling Basin (Fig S2), which covers most of south-east Australia, with the year effect estimates in virus prevalence in *Anatidae* (Fig 3B), and found a significant correlation (r = 0.782, N = 7, P < 0.04). Within the *Anatidae* and all-species model, for which we can assess both virus prevalence and seroprevalence, we found no correlation between the pattern of virus prevalence (r = -0.228, P = 0.623) and (r = 0.430, P = 0.215) seroprevalence across years. That is, we did not observe high virus prevalence in years of low seroprevalence. We also found that the year effects in seroprevalence are different between *Anatidae* and *Scolopacidae* (r = 0.621, P = 0.100).

### Role of eco-region in AIV prevalence

While the vast majority of our data set comprises samples collected in temperate Australia, 1950 samples were collected in arid Australia [largely *Anatidae*] and 1062 samples were collected in tropical Australia [largely *Scolopacidae*]. Interestingly, we only observed significant effects of eco-region on virus prevalence in *Anatidae* and on seroprevalence in the all-species taxon set. In the *Anatidae*, virus prevalence was higher in temperate regions as compared to arid regions (where percentages are calculated from the logit estimates depicted in Fig 3; 11.0%, 6.7-18.5).

## Discussion

Our holistic study of AIV evolutionary ecology in wild birds is unique for its use of a paired virological and serological data set. The inclusion of serological data expands the window of detectability of AIV. While AIV infections in individuals are only 7-11 days (Latorre-Margalef *et al*. 2014), anti-AIV antibodies may persist in the order of months to years (Tolf *et al*. 2013; Hill *et al*. 2016) and therefore population level seroprevalence is modulated by processes over much longer time scales like season, rainfall and migration (van Dijk *et al*. 2018). The use of serology allowed us to identify bird taxa that, while they are unlikely to be important reservoirs, are occasionally infected by AIV (*e*.*g*. Zebra Finch). Adding serology thus also adds credibility that the results of viral prevalence data are not influenced by missing prevalence peaks and non-representative sample collection. This would manifest as low AIV prevalence but higher seroprevalence (eg Silver Gull), although this could also be caused by exceptional long lived anti-AIV antibodies. Conversely, where AIV prevalence matches seroprevalence levels, this may suggest unbiased sampling. For instance, both low viral prevalence and seroprevalence in Sanderling and Wood Duck as compared to other related taxa, suggests that those results are not likely to be biased by our sample collection regime and that these two species are probably true outliers within these two AIV-reservoir species groups. Across all species combined and within the more traditional hosts (*Anseriformes* and *Charadriiformes*), analysing both serological data and virology data strengthened the interpretation of the various random and fixed effect explanatory variables, yielding largely overlapping and mutually supporting patterns.

While variation in susceptibility and competence among host species is a common feature of multi-host systems, the importance of phylogeny has rarely been explicitly addressed (Longdon *et al*. 2011; Greenberg *et al*. 2017; Barrow *et al*. 2019). Indeed, despite variation in prevalence across species in waterfowl or shorebird systems, the phylogenetic relationships among host species has never previously been integrated into statistical approaches to increase our understanding of the key drivers of AIV ecology. The strong phylogenetic effect found in the all-species comparison is unsurprising given the identification of *Anseriformes* and to a lesser extent *Charadriiformes* as highly competent AIV hosts compared to other bird taxa decades ago (Webster *et al*. 1992; Olsen *et al*. 2006). However, the finding of a strong phylogenetic effect within both the *Anatidae* and *Scolopacidae* is striking. That phylogeny is such an important covariate strongly suggests that genetic relatedness, perhaps including shared aspects of the immune response and/or virus susceptibility, are at play. The strong phylogenetic effects identified may also be key elements of host-virus co-evolution, and likely explain differences in host responses to infection, such as avoidance, resistance or tolerance.

Hosts and pathogens may co-evolve, with strong selective pressures acting upon both the host and the virus. It has long been argued that wild birds and AIV have undergone long term co-evolution, such that reservoir taxa may have adapted towards tolerance rather than resistance of AIV through mounting of a dampened immune response. In turn, (low pathogenic) AIV evolved low virulence in these hosts (Kuiken 2013). In this study we illustrate this co-evolution through a significant phylogenetic effect. Indeed, Longdon *et. al*. (2011) and Barrow *et. al*. (2019), who found phylogenetic signals in virus susceptibility in Drosophila and parasite susceptibility in birds, respectively, similarly argue that phylogenetic variation was driven by the generalised immune response and that there has likely been long term co-evolution between viruses/parasites and their hosts.

Beyond phylogenetic effects, species effects that are not driven by phylogeny also played an important. For instance, within the six species of the genus *Calidris* (*i*.*e*. Curlew Sandpiper to Red-necked Stint in Fig 2), we found large variation in prevalence. These marked species differences across closely related species could be due to variations in habitat preference. For example, Sanderling is the most marine and beach-dwelling of all *Calidris* species. In addition, among the *Anatidae*, the most distantly related species (the *Anas* ducks versus the Pink-eared Duck) had similar prevalence values, whereas the intermediated related Wood Duck had very low prevalence values. This is likely explained by foraging ecology, where Wood Duck is an exclusive grazing duck in contrast to the other species that dabble or filter feed.

This study provides the most comprehensive assessment of AIV ecology on the Australian continent. No previous studies have directly addressed host susceptibility, age or eco-region, while only two studies addressed year and season effects (Hansbro *et al*. 2010; Curran *et al*. 2014, 2015; Grillo *et al*. 2015; Ferenczi *et al*. 2016). First, in addition to species and phylogenetic effects, our statistical approach accounted for life history (age), seasonal, annual and environmental effects that are confirmed drivers of infection. As previously demonstrated, age is an important driver of AIV ecology. Higher AIV prevalence has been found in juvenile compared to adult ducks (Latorre-Margalef *et al*. 2014; van Dijk *et al*. 2014), and in Mute Swans the immune repertoire increases with age (Hill *et al*. 2016). Second, as noted previously, seasonal cycle is central to prevalence: prevalence peaks are associated with autumn migration in the temperate north (Latorre-Margalef *et al*. 2014; van Dijk *et al*. 2014), with the arrival of European migrants in Africa (Gaidet *et al*. 2012a; Gaidet 2016), and with rainfall in Australia, although for the latter this is often on a multi-year rather than annual scale (Ferenczi *et al*. 2016). A determinative feature of the southern hemisphere climate, particularly Australia, is the ENSO and IOD linked irregularity in both timing and location of wet and dry periods (Stuecker *et al*. 2017). As a result, breeding seasonality does not mirror that of northern hemisphere, rather some species may have elongated breeding times (5-7 months), or breeding may be competently opportunistic and take place at any time of the year with multiple broods in wet years (Halse & Jaensch 1989; Norman & Nicholls 1991). Therefore, with increased rainfall, more juvenile birds are recruited into populations, driving an increase in the proportion of immunology naïve birds in waterfowl populations, in turn modulating AIV prevalence (Ferenczi *et al*. 2016). Third, in addition to strong year effects associated with increased rainfall, we found that in the long-distance migratory *Scolopacidae* AIV prevalence was lowest just after their arrival from the breeding grounds and highest during the latter stages of the non-breeding season in Australia. Low population prevalence upon arrival may be due to parasites limiting migration (McNeil *et al*. 1994), and thus new-arrivals being preferentially free of pathogenic infections. Moreover, lower temperatures and lower UV levels during the latter stage of their Australian staging may be more conducive for virus survival (van Dijk *et al*. 2018). Finally, despite the sampling across the three eco-regions arid, tropical, and temperate being strongly biased towards the latter region, prevalence appeared lowest in arid environments, in line with the virus’ susceptibility to desiccation (Zarkov & Urumova 2013).

Taken together, in addition to confirming the role of climate as well as (ENSO-linked) rainfall and age on AIV prevalence, we provide new insights into AIV evolutionary ecology that define the specific processes that occur on the continent of Australia and expand our understanding of the factors that modulate AIV ecology across wild birds globally. Notably the strong phylogenetic and non-phylogenetic species effects revealed here highlight the importance of teasing apart these two overlooked factors in AIV ecology and evolution. Simultaneously considering the existence of strong phylogenetic and non-phylogenetic species effects, even within the two major AIV competent families, highlights how species-specific approaches are required in identifying reservoir communities, for understanding AIV dynamics in avian communities, and in evaluating spill-over risks of AIV from wildlife to livestock and humans.

## Acknowledgements

Collecting this large data set was only possible with the help of many. In particular we would like to mention the volunteers of the Victorian Wader Study Group and the Australasian Wader Study Group and in particularly Clive Minton, duck hunters from Geelong Field and Game and staff and students working with MK at Deakin University. We are grateful for the logistic support from Melbourne Water, Innamincka Station, Gidgealpa Station and Innamincka Regional Reserve. Finally, we thank Kim O’Riley and Malet Aban for contributing to sample screening.

